# Ethanol induces neuroimmune dysregulation and soluble TREM2 generation in a human iPSC neuron, astrocyte, microglia triculture model

**DOI:** 10.1101/2025.08.04.668469

**Authors:** Andrew J. Boreland, Yara Abbo, Xindi Li, Alessandro C. Stillitano, Siwei Zhang, Jubao Duan, Zhiping P. Pang, Ronald P. Hart

## Abstract

Alcohol use disorders (AUDs) affect substantial populations worldwide and increase the risk of developing cognitive impairments and alcohol-associated dementia. While chronic inflammatory signaling likely plays an important role in alcohol-associated neurological sequalae, the precise mechanisms underlying alcohol-associated neuropathology remain enigmatic. We hypothesize that alcohol leads to neuroimmune dysregulation among neurons, astrocytes, and microglia; and is perpetuated by innate immune signaling pathways involving cell-cell signaling. To investigate how alcohol dysregulates neuroimmune interactions in a human context, we constructed a triculture model comprising neurons, astrocytes, and microglia derived from human induced pluripotent stem cells. After exposure to ethanol, we observed significant differential gene expression relating to innate immune pathways, inflammation, and microglial activation. Microglial activation was confirmed with morphological analysis and expression of CD68, a lysosomal-associated membrane protein and marker for phagocytic microglial activation. A striking finding in our study was the elevation of TREM2 expression and, specifically, TREM2 alternative splice variants that are predicted to give rise to soluble TREM2. TREM2 has been reported to be a risk factor for Alzheimer’s disease. These results suggest that ethanol exposure in the brain may lead to increased microglial activation and production of soluble isoform named TREM2^219^ through alternate splicing. Deciphering the molecular and cellular mechanisms underpinning ethanol-related neuroimmune dysregulation within a human context promises to shed light on the etiology of AUD-related disorders, potentially contributing to the development of effective therapeutic strategies.

**Highlights:**

- We prepared a “triculture” of human iPSC-derived neurons, astrocytes and microglia
- Ethanol treatment produces substantial changes in gene expression with prominent effects on neuroimmune signaling
- Several microglia-specific genes are induced by ethanol in tricultures but not in cultures of microglia alone
- TREM2 expression is increased following ethanol treatment and results indicate a differential splicing of isoforms, encoding a soluble form of TREM2

## INTRODUCTION

Chronic neuroinflammation is a hallmark of alcohol use disorder (AUD)-related dementia^1,2^. Specifically, dysregulated neuroimmune interactions between neurons and glial cells, such as astrocytes and microglia, present a candidate mechanism for understanding how these disorders develop. We now know that microglia, the brain’s primary immune effector cells, play a prominent role in healthy brain function and, when dysregulated, are directly implicated in cognitive disease related pathology^3–5^. Therefore, unraveling the molecular and cellular mechanisms underlying human neuroimmune dysregulation, in relation to ethanol, one of the most prolific and widely consumed substances worldwide, may lead to new insights crucial for understanding and treating alcohol use disorders and neurological sequelae.

Despite extensive studies investigating the consequences of ethanol exposure in rodent models^6–9,10^, limited data exists on ethanol’s effects on human brain cells including neurons, astrocytes, and microglia. This is particularly important since many differences exist between rodent and human brain development^11^ and noncoding, regulatory genetic elements are often species-specific^12^. Dysregulated neuroimmune interactions between neurons and glial cells, such as astrocytes and microglia, may underlie aspects of alcohol-related neuropathology. Human-specific neuroimmune interactions, however, have been historically difficult to study, partly due to a paucity of human brain tissue for research. Human induced pluripotent stem cell (iPSC)-derived neural models provide highly manipulatable model systems which have been shown to model neuroinflammation^13^ including several studies from our lab^14–18^.

One sign of chronic inflammation in the brain is *TREM2*, encoding the triggering receptor on myeloid cells (TREM2) protein. Importantly, *TREM2* is a risk gene for Alzheimer’s Disease (AD) with rare, heterozygous variants, such as *TREM2* R47H, being significantly associated with increased risk for not only AD^19^, but also Parkinson’s Disease (PD)^20^, and amyotrophic lateral sclerosis (ALS)^21,22^. Activation of TREM2 not only modulates microglial activation states but is also crucial for the acquisition of a ’disease-associated microglia’ (DAM) profile^23^. TREM2 is essential for microglial fitness during stress events^23^, such as an individual ingesting alcoholic beverage and the ensuing cellular stress [reviewed in^24,25^]. Importantly, due to the affinity of this receptor for lipid-associated complexes such as myelin and beta amyloid, TREM2 is believed to play an important role in AD onset and progression^26^. TREM2 can also exist in a soluble form, sTREM2, which has been associated with amyloid-beta plaques in AD and shows elevated levels in AD patient cerebral spinal fluid over the course of the disease^27^.

We hypothesize that ethanol leads to neuroimmune dysregulation among neurons, astrocytes, and microglia, and is perpetuated by innate immune signaling pathways, likely involving cell-cell signaling. Here, we constructed a triculture model comprising iPSC-derived neurons, astrocytes, and microglia to investigate how ethanol dysregulates neuroimmune interactions in a human-specific context.

## RESULTS

### Generation of a human iPSC-derived neural triculture

Adapting previous protocols^28^, we prepared neural progenitor cells (NPCs) from iPSC using neural induction by dual-SMAD inhibition (**Figure 1A**). iPSC-derived microglia, which have a mesodermal yolk-sac ontology, were generated separately using previously published protocols^17,28,29^ (**Figure 1A**). To form the triculture, NPCs were cultured in neuronal maturation conditions for 7 weeks to allow neurogenesis and astrogenesis before microglia were added. Cell identities in the triculture were confirmed by immunofluorescence staining for MAP2 (microtubule-associated protein 2) to label neurons, GFAP (glial fibrillary acidic protein) to label astrocytes, and IBA1 (ionized calcium-binding adapter molecule 1) to label microglia (**Figure 1B**). Quantification of immunostained cultures indicates a typical proportion of 35% neurons, 52% astrocytes and 13% microglia. Microglia were found to integrate into the culture and interact with neuronal dendrites, suggesting physiological surveillance activity^15,28^ . Taken together, these results indicate formation of an iPSC-derived triculture model system for evaluating the effects of ethanol on neuroimmune cell-cell interactions among neurons, astrocytes, and microglia.

**Figure 1:**
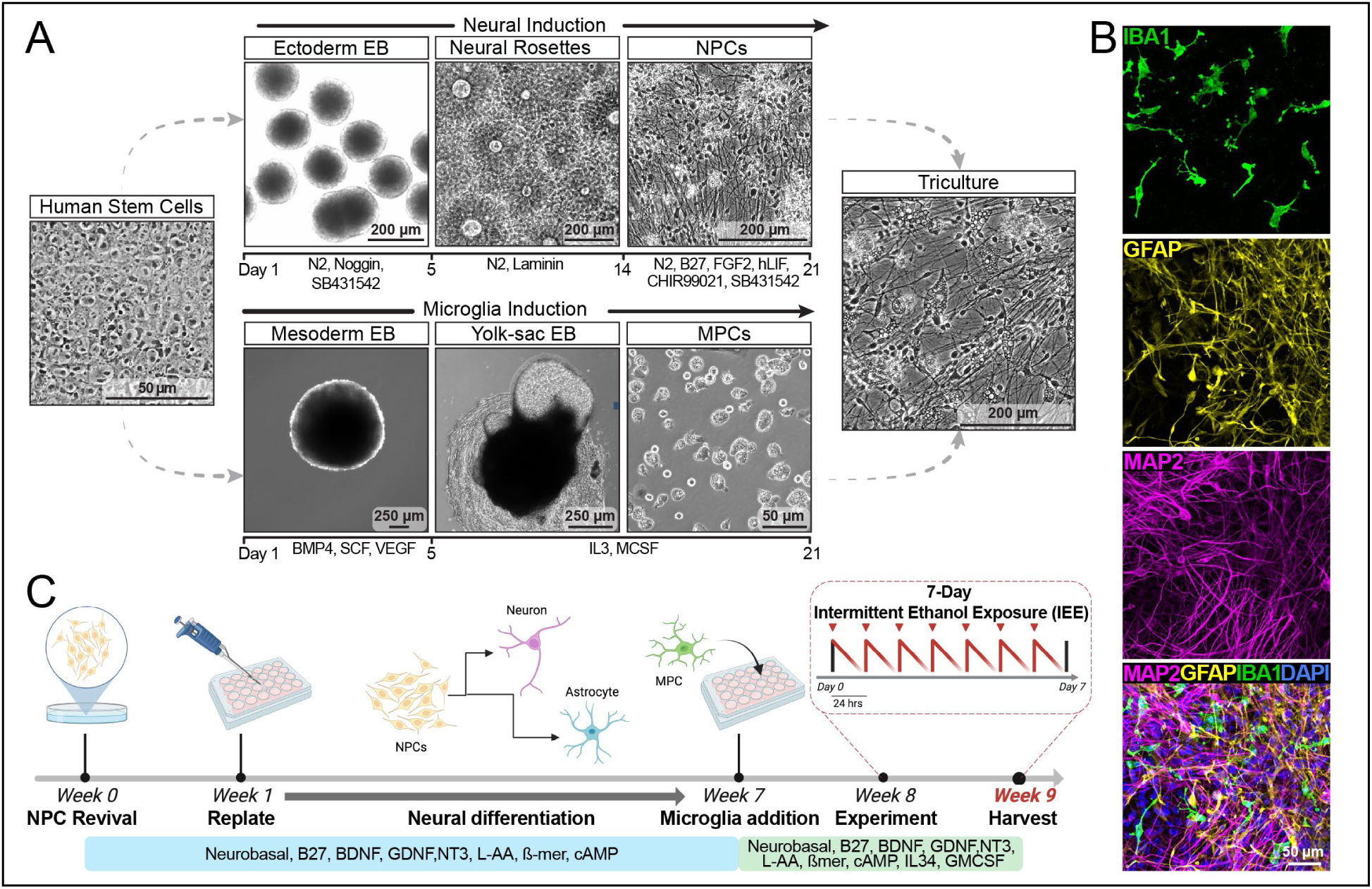
Generation of a human stem cell derived neuron, astrocyte, and microglia triculture. (A) A schematic procedure for preparing tricultures from iPSC-derived neural progenitor cells (NPCs) and microglial precursor cells (MPCs). See Methods for details. (B) Representative confocal images of immunohistochemical-stained tricultures showing microglia marker IBA, astrocyte marker GFAP, neuronal dendrite marker MAP2 and an overlay combining all markers. (C) Graphical summary of the experimental procedures. After microglia are added to neurons and astrocytes prepared from NPCs (week 7), cultures were treated with ethanol using an IEE daily replenishment schedule for 7 days, with concentrations dropping each day, presumably due to evaporation from the cultures (see cartoon of ethanol concentrations).

### Intermittent ethanol exposure (IEE) induces diverse inflammation-associated differential gene expression

To model the consequences of frequent alcohol drinking on the CNS, we employed an IEE paradigm^18,30^ on tricultures (**Figure 1E**). Ethanol was applied once every 24 hours to reach peak concentrations of 0 mM, 20 mM, 40 mM, or 75 mM for 7 days to mimic a repeated daily drinking pattern seen in individuals with AUD^18,30^. Importantly these ethanol concentrations correlate to physiological blood alcohol concentrations (BAC) with increasing clinical significance and impairment^31^; a 20 mM ethanol concentration corresponds to a peak of 0.092% BAC, 40 mM corresponds to peak of 0.184% BAC, and 75 mM ethanol concentration corresponds to a peak of 0.346% BAC, corresponding to severe intoxication^32^. In tissue culture, there is a rapid decrease in concentration likely to be primarily due to evaporation **(Figure 1C)**. Using half-life data from our previous study^30^, we calculated the mean ethanol concentration over a 24 hour period to be 10.26 mM for 20 mM peak concentration, 18.88 mM for 40 mM peak concentration, and 28.79 mM for 75 mM peak concentration.

To identify ethanol-regulated gene expression, we assayed tricultures using RNA sequencing. Principal component analysis of the top 500 most variable genes indicated that the primary distinction among samples was ethanol concentration (**Figure 2A**). Results revealed 189 upregulated and 88 downregulated differentially expressed genes (DEG) comparing 20 mM ethanol to control, 557 upregulated and 278 downregulated genes comparing 40 mM ethanol to control, and 804 upregulated and 462 downregulated genes comparing 75 mM ethanol to control (**Figure 2B**). Interestingly, a culture containing only microglia treated with a similar 7-day IEE, had only 185 upregulated but 516 downregulated genes comparing 75 mM to control (**Supplemental Figure 2A**). This divergence may represent results from the combination of cell types in the triculture or it may reflect a more complex cellular and gene regulatory environment after ethanol in the multiple cell-type context, including cell-cell interactions among neurons, astrocytes, and microglia. Indeed, comparing the up and downregulated DEG between the triculture and monoculture revealed relatively little overlap (**Supplemental Figure 2B**). Differential gene expression analysis highlighted significant upregulation of innate immune related genes by ethanol, including TREM2, TLR2, TLR4, CCL3, and IL10 in all doses with increasing effect at increased dose (**Figure 2C**). We then conducted Reactome pathway analysis and found significant enrichment for terms relating to positive regulation of TNFα, interleukin-6 production, inflammatory response, response to pathogens, and DNA-damage response (**Figure 2D**).

**Figure 2:**
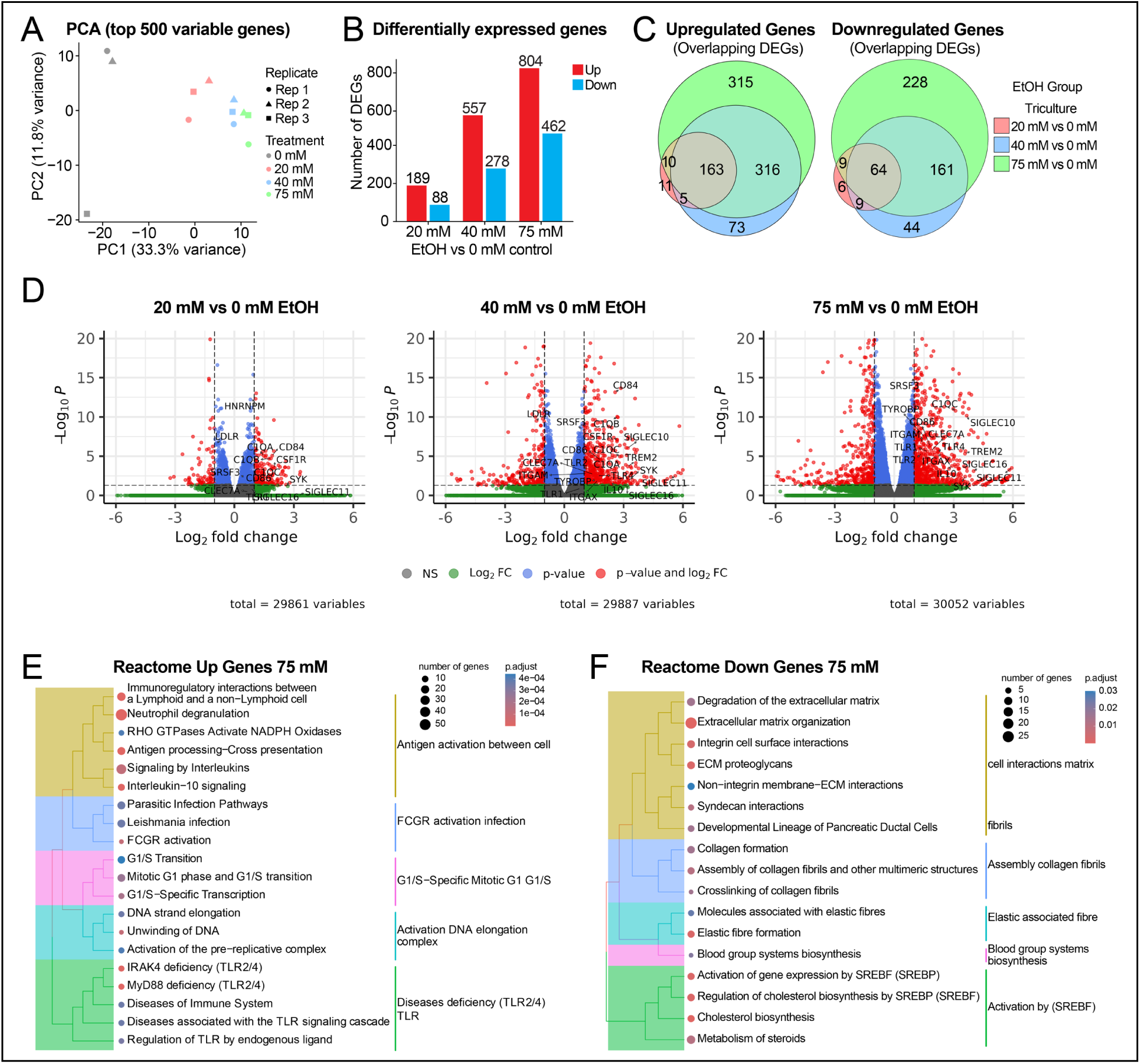
Intermittent ethanol exposure induces robust inflammation associated differential gene expression. (A) Principal component analysis of RNAseq samples from 7-day IEE. n = 3 independent experiments. (B) Differentially expressed gene (DEG) counts for 7-day CIE tri-culture RNAseq data. Volcano plots depicting DEG. (D) Reactome gene ontology analysis. (E) Reactome disease ontology analysis.

To identify enriched pathways from ethanol-regulated genes, we conducted gene set enrichment analysis (GSEA). As shown in **Figure 3A**, ethanol results in upregulation of biological processes relating to DNA damage response such as DNA repair and DNA replication, consistent with studies demonstrating DNA damage induced by ethanol and its metabolites^18,33,34^. Analysis of the cellular component terms revealed enrichment of pathways relating to the chromosomal region, vacuolar membrane, and lysosomal membrane across ethanol concentrations (**Figure 3B**). Finally, analysis of the molecular function ontology showed enrichment for phospholipid binding, histone binding, and mRNA binding (**Figure 3C**).

**Figure 3:**
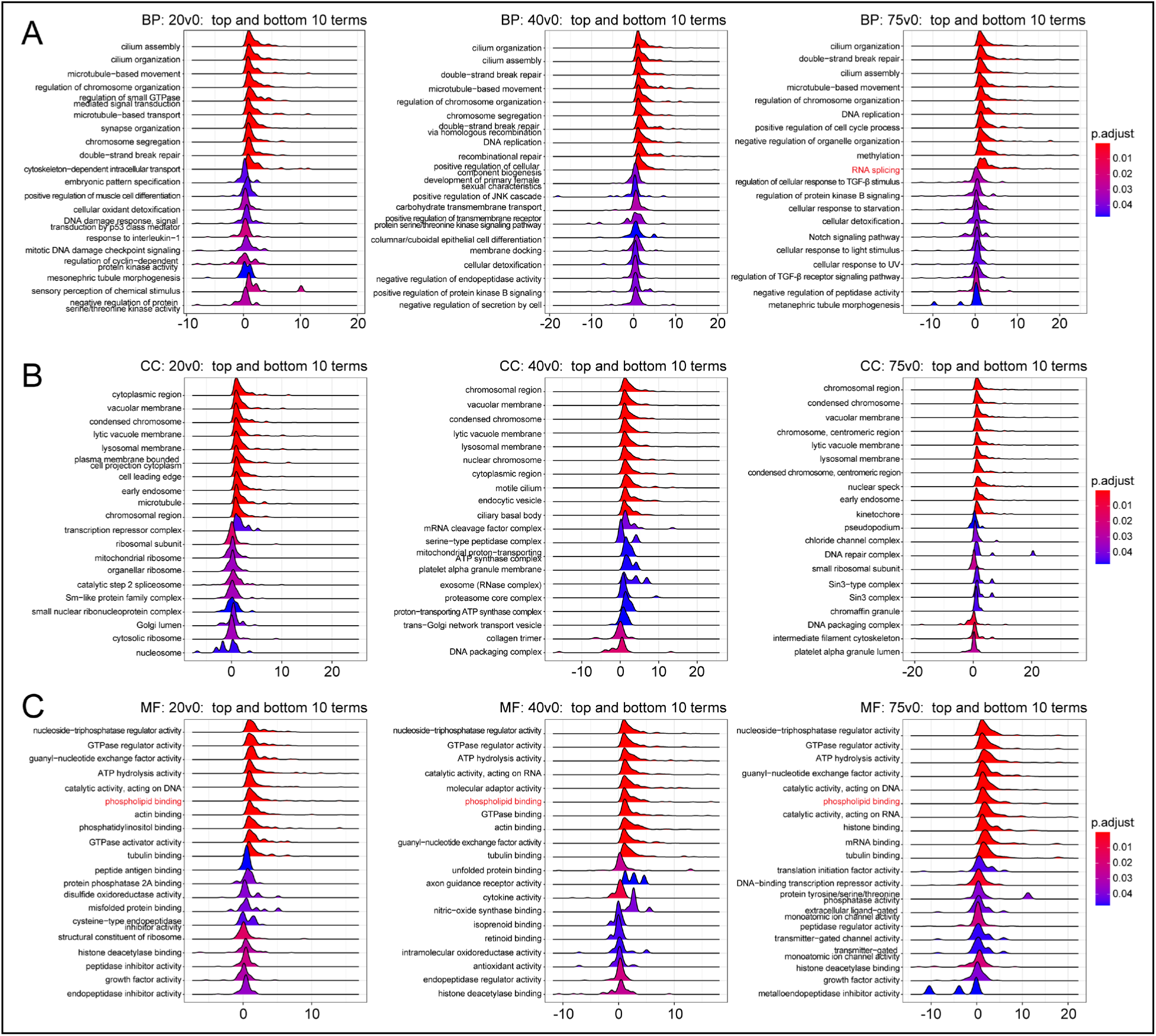
Gene set enrichment analysis (GSEA) reveals gene pathways altered by ethanol exposure. (A) Gene ontology Biological Processes comparisons between 20 mM vs 0 mM, 40 mM vs 0 mM, and 75 mM vs 0 mM ethanol exposed tricultures at Day 7. (B) Gene ontology Cellular Component comparisons between 20 mM vs 0 mM, 40 mM vs 0 mM, and 75 mM vs 0 mM ethanol exposed tricultures at Day 7. (C) Gene ontology Molecular Function comparisons between 20 mM vs 0 mM, 40 mM vs 0 mM, and 75 mM vs 0 mM ethanol exposed tricultures at Day 7.

### Ethanol induces microglial activation in a human neural triculture model

Since the RNA sequencing data suggested activation of several inflammatory pathways, including many genes with expression normally restricted to microglia, we sought to further characterize ethanol-associated microglia activation in the triculture. To confirm increased expression of microglia-specific genes associated with inflammatory activation following the 7-day IEE paradigm, we used reverse transcription/quantitative PCR (RT-qPCR) to examine the expression of several genes associated with microglial activation and inflammatory response. IBA1 is a calcium binding adapter important in microglia membrane ruffling and motility^35^. Consistent with the RNAseq data, we observed significantly increased IBA1 expression at 40 mM and 75 mM ethanol **(Figure 4A)**. We then assessed expression of two other genes related to microglial activation, M-CSF (macrophage colony stimulating factor) and CD86 (Cluster of differentiation 86), a canonical marker for M1 activated microglia^36^. We found that ethanol significantly increased M-CSF expression at all doses **(Figure 4A)**. CD86 displayed increased expression in all conditions compared to control, albeit significantly only at 40 mM **(Figure 4A)**.

**Figure 4:**
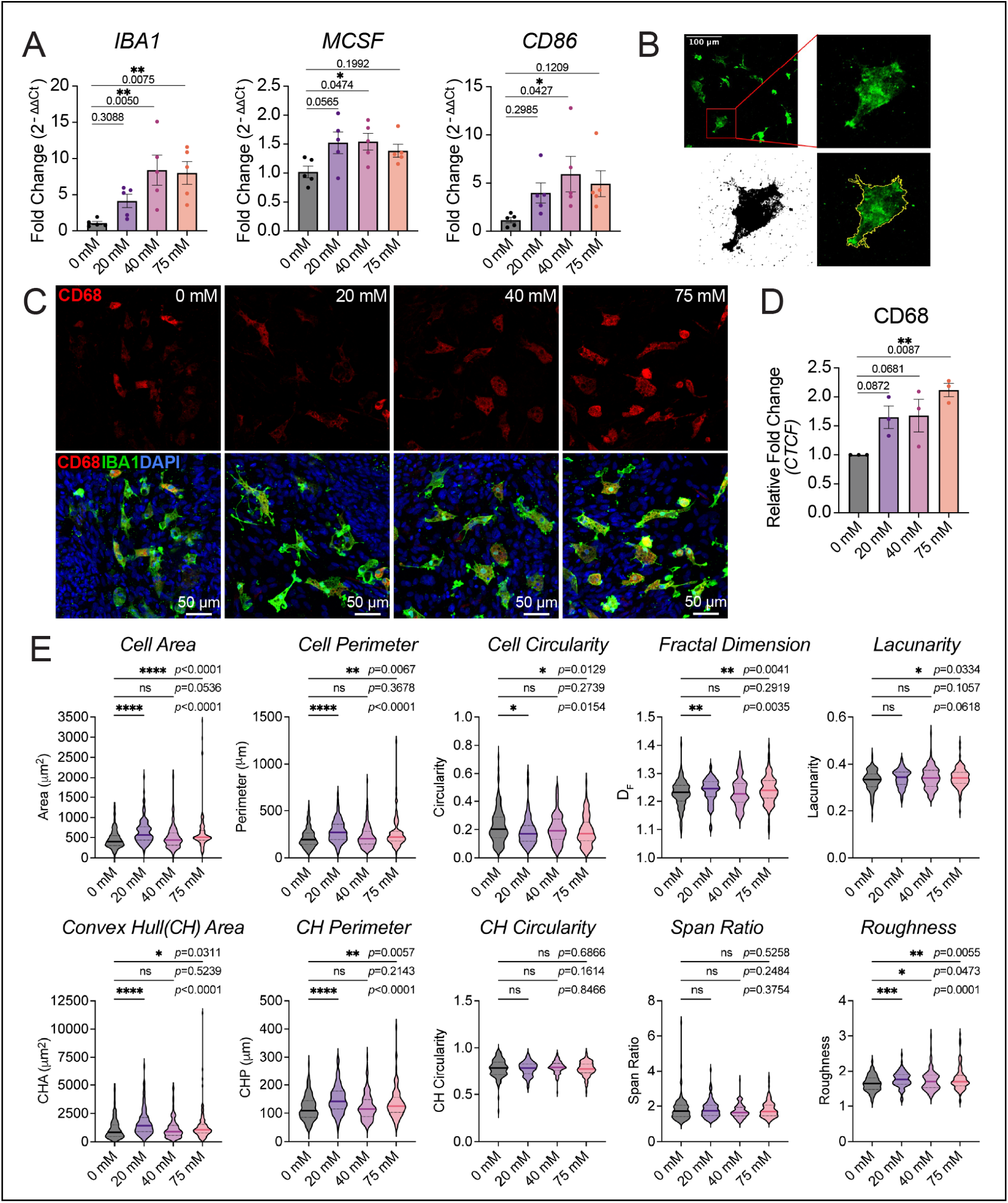
Ethanol produces microglial activation and morphological changes. (A) RT‒qPCR analysis of IBA1, MCSF, and CD86 mRNAs from 7-day IEE neural tri-culture, n = 5 independent experiments, one-way ANOVA, Dunnett’s multiple comparisons test. Data are presented as mean ± SEM. (B) Representative images showing how ROI are made from mask of IBA1 channel. A mask is constructed from the IBA1 signal to outline the cell of interest. This mask is applied to a separate channel to measure fluorescence signal for a second antibody. (C) Representative images of CD68 immunostaining overlying IBA1 in all ethanol doses. (D) Quantification of CD68 Corrected Total Cell Fluorescence (CTCF) normalized to each control (0 mM) replicate. n = 3 independent experiments, one-way ANOVA * P = 0.0151, Holm-Šídák’s multiple comparisons test. Data are presented as mean ± SEM. (E) Morphological analysis of microglia binary outlines using *FracLac ImageJ*. n = Cells/Batch: 340/3 for 0 mM, 299/3 for 20 mM, 319/3 for 40 mM, and 325/3 for 75 mM. Kruskal-Wallis Test, Dunn’s multiple comparison test. Data are presented as mean ± SEM.

To further investigate how ethanol exposure affects microglial activation, we quantified protein expression using immunohistochemistry against CD68, a transmembrane protein localized on lysosomes and strongly upregulated with inflammation^36^. CD68 is widely considered to be one marker for phagocytic microglial activation^36^. Regions of interest (ROIs) were made for each microglial cell based on IBA1 staining and cell segmentation, which allowed for individual microglial-specific quantification of CD68 fluorescence **(Figure 4B)**. CD68 staining and quantification revealed increased expression trending towards significance at 20 and 40 mM and significantly elevated expression at 75 mM **(Figure 4C,D).** Results indicate that levels of CD68 increase in IBA^+^ cells, consistent with microglial activation.

Microglial morphology is closely related to function and activation states. To assess ethanol’s effects on microglial morphology, we used FracLac^37^ analysis to quantify various morphological parameters on a single cell level using cell-specific ROIs generated from segmentation of IBA1 staining. First, we quantified parameters for cell size including total cell area (µm^2^) and perimeter (µm). Ethanol exposure led to significant increases in cell size and perimeter in 20 and 75 mM, but not 40 mM **(Figure 4E)**. Second, we assessed parameters for cell spread including convex hull (CH) area and CH perimeter; CH was calculated using the smallest convex polygon that can enclose the outline of a given cell. In accordance with cell area and perimeter increasing after ethanol, CH area and perimeter also increase in 20 and 75 mM, but not 40 mM. Third, we quantified parameters for cell shape including circularity and span ratio. Circularity, assessed for both cell and convex hull, measures the degree to which the cell or convex hull approaches a perfect circle (the circularity of a circle is 1)^38^ **(Fig. 4E)**. While cell circularity slightly decreased with ethanol, no change was observed for CH circularity. Likewise, span ratio showed no changes with ethanol exposure. Fourth, we quantified parameters relating to cellular complexity, heterogeneity, including fractal dimension, lacunarity, and roughness. Fractal dimension (D_B_) measures how image details change with increasing scale such that for two-dimensional images, cell complexity increases as D_B_ approaches 2^37^. We found ethanol causes subtle increases in fractal dimension with significant changes observed in 20 mM and 75 mM conditions. Lacunarity, which describes the heterogeneity/rotational and translational invariance of the cell shape/outline ^38^, showed subtle changes across conditions and only reached significant differences at 75 mM ethanol. Finally, cell roughness describes the ratio of the cell perimeter over the convex hull perimeter ^38^. We found that ethanol caused significant changes in cell roughness across 20 mM, 40 mM, and 75 mM concentrations. Together with the RT-qPCR and immunocytochemistry results, the morphological changes support the conclusion that ethanol exposure leads to microglial activation.

### Ethanol primes the NLRP3 inflammasome in triculture

NLRP3 (nod-like receptor family pyrin domain containing 3) expression has been identified in all major cell types of the brain, including neurons^39^, astrocytes, oligodendrocytes, and microglia [Reviewed in^40–42^]. In an earlier study, we demonstrated ethanol-associated priming of the NLRP3 inflammasome pathway in human iPSC and NPC cultures^43^. To assess ethanol-associated priming of the NLRP3 inflammasome pathway in triculture conditions, we first quantified NLRP3 RNA expression from bulk RNA using RT-qPCR. As shown in **Figure 5A**, NLRP3 RNA expression increased ∼5-fold following 20 mM IEE and significantly increased ∼8-9 fold with both 40 mM and 75 mM IEE. Next, we used immunocytochemical staining and confocal imaging to quantify NLRP3 protein expression after IEE. We found NLRP3 protein expression increased with each ethanol dose and significantly increased with 75 mM ethanol (**Figure 5B,C**). We then examined expression of Caspase-1 (CASP1), the component of the inflammasome complex that cleaves and activates pro-inflammatory IL1ß, by immunocytochemistry. RT-qPCR revealed significantly increased IL1ß mRNA expression from 40 mM and 75 mM ethanol exposure (**Figure 5D**), suggesting increased production of pro-IL1ß/IL1ß. 75 mM ethanol appears to cause increased protein expression of IL1ß, although mostly localized to microglia-like structures (**Figure 5E**). CASP1 expression also appears to increase but with expression more widespread and less restricted to microglia-like structures (**Figure 5E**), suggesting the presence of cell-cell signaling following activation of microglia. Together these data suggest priming of the NLRP3 inflammasome pathway in triculture conditions following 20 mM, 40 mM, and 75 mM IEE.

**Figure 5:**
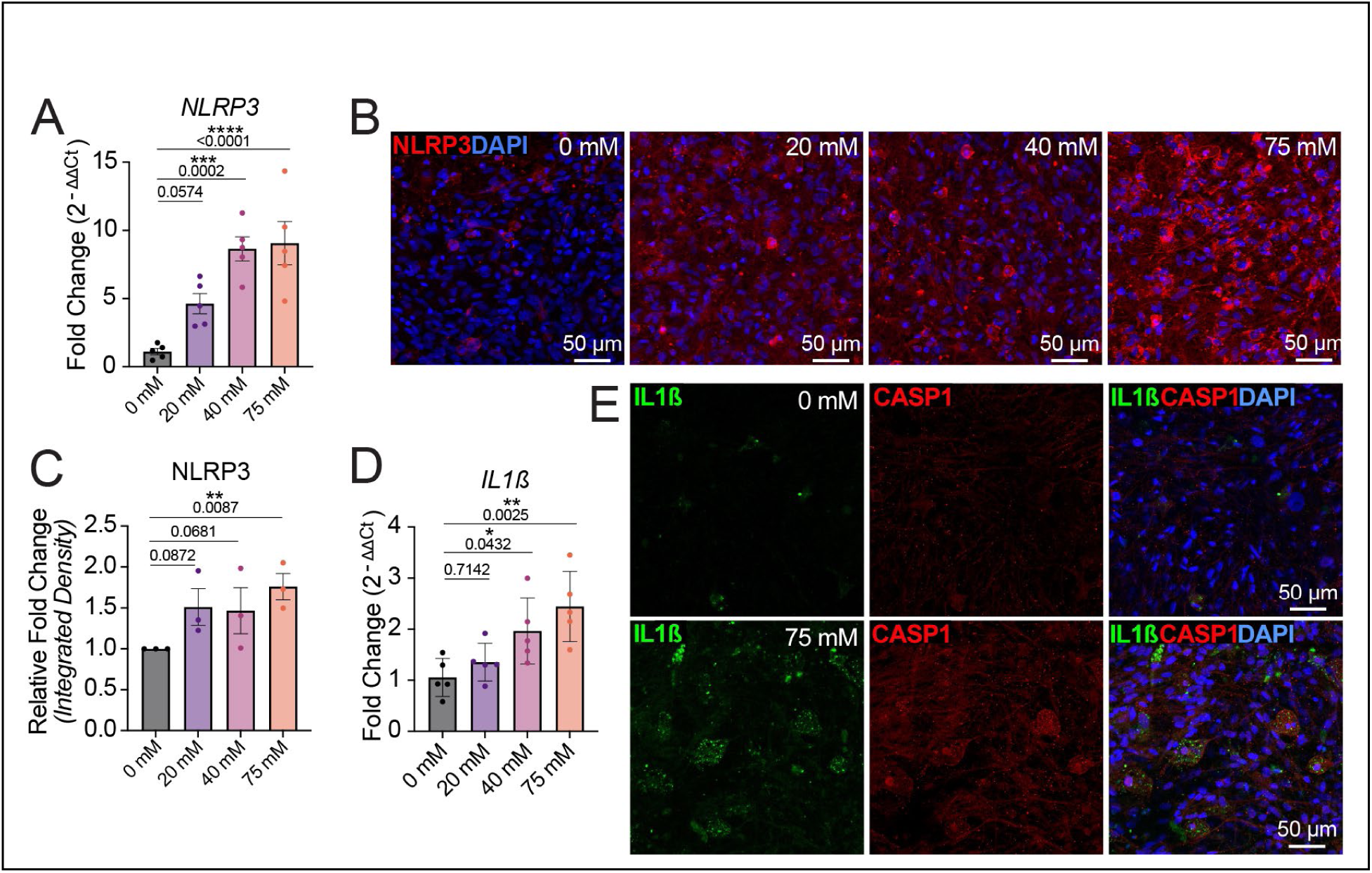
Dose dependent activation of the NLRP3 inflammasome pathway after ethanol exposure. (A) qPCR analysis of NLRP3 mRNA from 7-day IEE neural tri-culture, n = 5 independent experiment. (B) Representative images of NLRP3 expression in a 7-day IEE tri-culture with identical parameter. (C) Quantification of NLRP3 fluorescence integrated density normalized to each control (0 mM) replicate. n = 3 independent experiments. (D) qPCR analysis of IL1ß mRNA from 7-day IEE neural tri-culture, n = 5 independent experiment. (E) Representative images of IL1β and CASP1 expression in a 7-day IEE tri-culture with identical parameters.

### Ethanol exposure alters TREM2 splicing and elevates sTREM2

Among the ethanol-upregulated genes in the RNAseq results is TREM2. *TREM2* encodes multiple mRNA and protein isoforms (**Figure 6A**), producing a soluble form (sTREM2) by either enzymatic cleavage and shedding of the TREM2 receptor ectodomain, or alternate splicing of TREM2 transcripts (**Figure 6B**). While TREM2^230^, the predominant isoform, encodes the full-length 230 amino acid protein, two other transcript variants are known to exist, namely TREM2^222^ and TREM2^219^, both of which give rise to soluble species due to the missing transmembrane domains. Transcript isoform analysis of ethanol-exposed tricultures revealed that 72.44% were full-length TREM2^230^ isoform, 27.56% alternately spliced TREM2^219^ isoform, and no detection of TREM2^222^ at the 75 mM ethanol dose (**Figure 6C**). Comparing 0mM, 20mM, 40mM, and 75 mM IEE, we found both 40 mM and 75 mM induced significant increases in the TREM2^230^ isoform (**Figure 6C**). Importantly, all doses of ethanol caused a significant increase in TREM2^219^ transcript levels (**Figure 6C**). While full-length TREM2^230^ is detected even under control conditions, TREM2^219^ transcripts were barely detectable (**Figure 6D**). To validate these findings, we performed RT-qPCR on RNA from ethanol-treated tricultures using primers specific for each transcript variant^44^. Ethanol increased expression of TREM2^230^ at each dose with significant increases at 75 mM, no changes in TREM2^222^, and significant increases in TREM2^219^ at 75 mM (**Figure 6E**). Together, these data suggest that ethanol exposure increases TREM2 expression and generation of sTREM2 species in the brain via increases in the alternatively spliced TREM2^219^ transcript variant.

**Figure 6:**
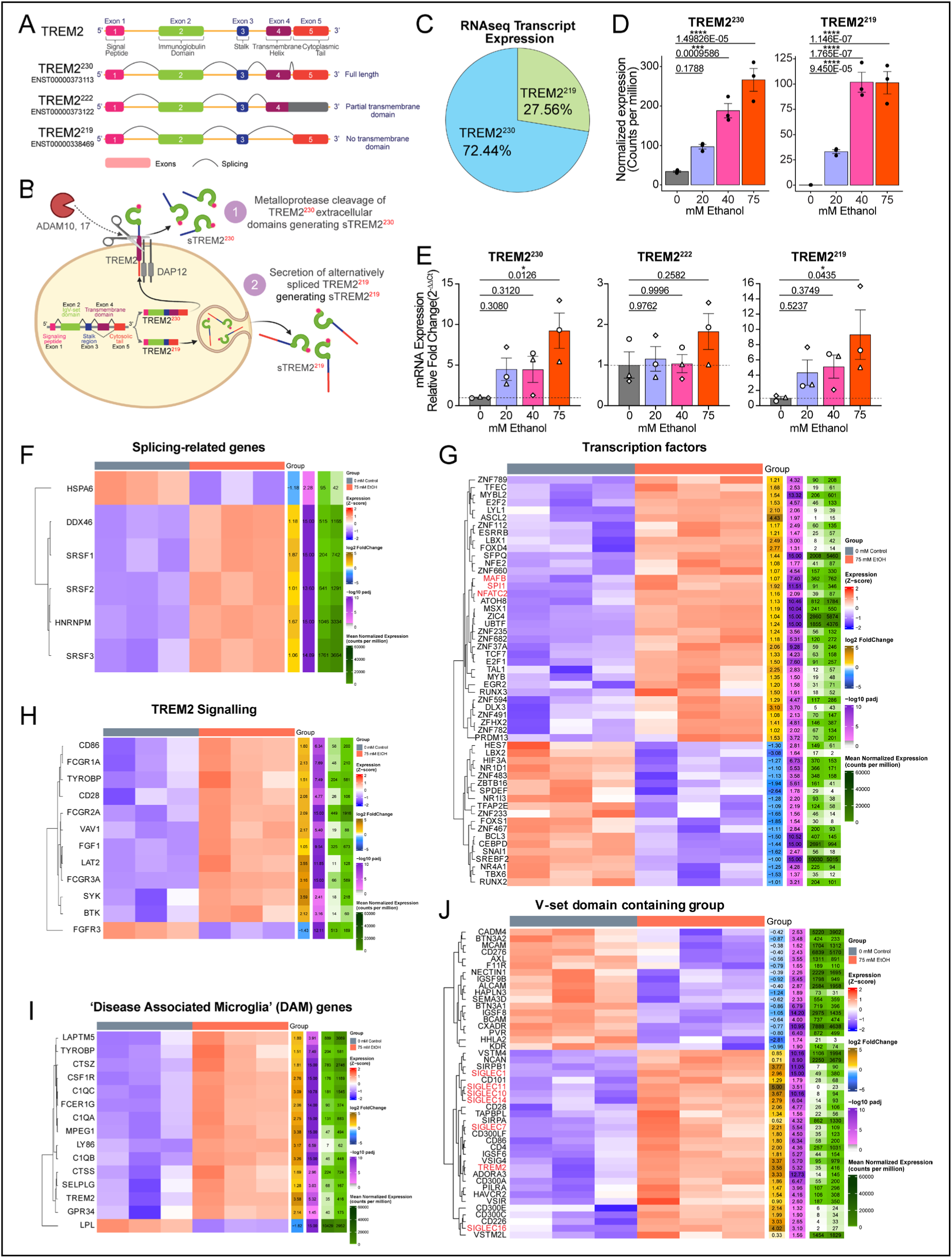
EtOH increases TREM2 and sTREM2 isoform expression. (A) Schematic showing the three splice isoforms of TREM2 from Ensembl. (B) Diagram showing multiple mechanisms of sTREM2 generation from TREM^230^ and TREM^219^ transcript variants (C) Proportion of TREM2 splice variants after 7 days of 75 mM IEE. (D) TREM2^230^ and TREM2^219^ transcript expression after 7 days of IEE in 75 mM condition. (E) RT-qPCR analysis of TREM2 splice variant isoforms in a 7-day IEE treated neural tri-culture. (F) A heatmap depicting significantly expressed splicing-related genes. (G) A heatmap depicting significantly expressed transcription factor genes. (H) A heatmap depicting significantly expressed TREM2 signaling genes. (I) A heatmap depicting significantly expressed DAM genes with TREM2-dependent expression. (J) A heatmap depicting significantly expressed V-set immunoglobulin domain genes.

Regulation of TREM2 could involve multiple mechanisms. Ethanol has been found to affect differential splicing of RNA transcripts and regulation of selected splicing factors in neurons^45–51^. We examined the regulation of several common genetic splicing factors and found significant differences in multiple splicing-related genes following 75mM IEE ethanol treatment (**Figure 6F**), including increases in DDX46, SRSF1, SRSF2, HNRNPM and SRSF3. One factor, HSPA6, was downregulated by ethanol. Most are similarly regulated with lower ethanol doses (not shown). While it is unknown which mechanisms specifically alter TREM2 splicing patterns, it is clear that ethanol has the potential to regulate mechanisms that could explain the differential splicing of TREM2. Many transcription factors are also affected by 75 mM IEE ethanol treatment (**Figure 6G**). A large up-regulated cluster includes MAFB and SPI1, both of which have been implicated in TREM2 gene regulation and are known to regulate a broad array of inflammatory genes^52,53^. NFATC2, aka NFAT1, a known modulator of TLR4-associated microglial activation^54–56^, was also upregulated and clustered with MAFB and SPI1^52,53^. Furthermore, a group of proteins has been identified as binding partners or signaling intermediates of TREM2^57,58^. Most of these are similarly up-regulated following 75 mM IEE ethanol treatment (**Figure 6H**), potentially providing enhanced opportunity for up-regulated and/or alternatively spliced TREM2 to signal additional mechanisms^44,58,59^. Expression patterns also demonstrate the activation of many disease-associated microglia (“DAM”) genes **(Figure 6I**), agreeing with previous results (**Figure 4**), which have previously been shown to have TREM2-dependence^53,60,61^. Finally, TREM2 is a member of a family of genes, including several sialic acid-binding immunoglobulin-like lectin (SIGLEC) family members, that share an immunoglobulin V-set domain^62,63^. The V-set domain is part of a larger immunoglobulin-like superfamily crucial for cell-surface recognition and cell-cell interactions^64,65^. While some V-set domain genes are down-regulated, TREM2 clusters with SIGLEC1, 11, 10, 14, and 16 in the up-regulated group (**Figure 6J**). Ethanol regulation of TREM2, along with a broad array of intercellular signaling components, suggests activation of a complex immunomodulatory network within the triculture.

## DISCUSSION

In this study, we used a human stem cell-derived neural triculture of neurons, astrocytes, and microglia to evaluate ethanol’s effects on neuroimmune pathways, microglial activation, and interactions among these three cell types. Until recently, evidence of neuroimmune modulation by ethanol has largely been restricted to rodent studies, partly due to a lack of primary human brain tissue for research. After IEE exposure, we observed significant differential gene expression in many genes relating to innate immune pathways, inflammation, and microglial activation. Notably, we identified not only upregulation of TREM2 mRNA, but a shift in isoform composition, suggesting an increase in sTREM2 production, which would trigger additional pathways in co-cultured cell types.

Ethanol induces an increase in inflammatory markers, including IBA1, MCSF, CD86, and CD68 (**Figure 4**). These genes are primarily associated with microglia; immunocytochemical detection, restricted to IBA1^+^ microglia, demonstrated that these markers increased in microglia as expected. Furthermore, since microglial morphology is tightly linked with inflammatory status, we identified changes in several cellular morphology parameters following ethanol treatment, including cell area, shape, and perimeter complexity. All of these markers were previously found to be induced by ethanol in microglial monocultures^18^. Both of these studies, however, relied on bulk RNA sequencing, which did not allow direct analysis of cell type-specific gene expression. However, several microglia-specific genes associated with inflammatory activation were induced by ethanol in tricultures but not in microglial monocultures (**Supplemental Figure 2**), suggesting that cell-cell interactions may amplify and/or extend microglial activation^66^.

In addition to the transcription data indicating microglial activation, we also found upregulation of CD68, a lysosomal-associated membrane protein and marker for phagocytic microglial activation^36^. This suggests that ethanol may modulate lysosomal activity and thus may have implications in microglial phagocytic functions and waste clearance. Interestingly, CD68 has been found to be upregulated in post-mortem studies of brains from individuals with AD^67^. In contrast, CD68 expression appears more varied in rodents following ethanol. For example, no changes in CD68 protein expression were observed in a 4-day binge alcohol model in rats^68^, while in a 5-month chronic ethanol mouse model, CD68 protein expression was found to increase^69^. While these differences may be due to dosage and length of exposure, an additional, prolonged 6-week ethanol paradigm in mice found decreased CD68 expression after ethanol exposure^70^. Such varied results in the literature suggest ethanol’s effects on CD68 expression and consequent microglial activation are quite complex; various factors including the dosage, duration, and frequency of ethanol exposure in addition to species differences and heterogenous genetic backgrounds likely contribute to the discrepancies found within the literature.

Previously, our group showed that ethanol-treated iPSCs and NPCs exhibited activation of the NLRP3 inflammasome pathway^43^. Inflammasomes are innate immune sensors that induce inflammation and ultimately pyroptosis^71^; importantly, aberrant and chronic activation of inflammasomes has been linked to various neurodegenerative diseases including multiple sclerosis and AD^71^. Indeed, in the triculture model, we also observe activation of the NLRP3 inflammasome as supported by increased gene and protein expression of NLRP3 and associated NLRP3 inflammasome complex proteins CASP1 and IL1ß (**Figure 5)**. Results suggest that ethanol likely leads to priming and activation of the NLRP3 inflammasome in multiple cell types. Importantly, a functional NLRP3 inflammasome should facilitate pro-IL1ß processing to IL1ß and subsequent secretion. While IL1ß is vital for host defense against pathogens, it can also exacerbate damage in other chronic disease states. An extremely potent cytokine, IL1ß has well-studied roles in neuropathic pain, inflammation, and autoimmunity; interestingly, evidence also suggests a role for IL1ß as a neuromodulator and central player in neuronal-glial interactions^72,73^.

Compared to human microglia-only cultures, when co-cultured with neurons and astrocytes, microglia exhibit an enhanced or altered response to ethanol (**Supplemental Figure 2**). In addition to finding surprisingly few genes up- or down-regulated in common between microglia and tricultures by 75 mM IEE (many of which are undoubtably due to expression in cell types other than microglia), there are several microglial-specific genes uniquely induced in tricultures (**Supplemental Figure 2C**). This includes several members of the SIGLEC family, which encompasses a group of transmembrane immunoglobulin-type lectins that bind sialic acid moieties on cell surface glycoproteins and glycolipids [Reviewed in^74^]. Interactions with SIGLECs play crucial roles in cell-cell communication, cell-pathogen recognition, and immune modulation. The SIGLEC family has undergone rapid evolution^63,75^ including the loss of certain family members (SIGLEC13 and SIGLEC17)^76^ and the rise of other distinct hominid-specific paralogs (SIGLEC11 and SIGELC16)^76,77^, demonstrating why it is important to study these pathways in human-specific models. In one example, SIGLEC11 is an inhibitory SIGLEC that recognizes sialyation of the neuronal glycocalyx, inhibits microglial phagocytosis, and has been shown to alleviate neurotoxicity^78^. This is countered by SIGLEC16, which has an activating role facilitated by an intracellular domain that can associate with DAP12 and potentiate TREM2-like signaling^77^. In addition to the SIGLEC receptor, the sialyation state of the neuronal glycocalyx can impact SIGLEC responses [Reviewed in^79^]. For instance, desialyation around synapses and subsequent accumulation of C1q protein at these locations initiates microglial trogocytosis or synaptic pruning^80^.

A striking finding of our study was the dramatic elevation of TREM2 mRNA expression and alternative splicing following the IEE paradigm. These results agree well with GSEA predictions, which found enrichment of biological process term “RNA splicing” (**Figure 3A**) and molecular function term “phospholipid binding” (**Figure 3C**). In contrast, most other models where TREM2 mRNA is regulated show down-regulation or loss of function^81–85^. Future studies should explore the relevance of increased TREM2 expression on neuronal viability and synaptic transmission, since earlier work largely focused on loss of function variants such as the R47H polymorphism^86,87^. Interestingly, studies on the normal function of TREM2 have shown a neuroprotective effect. For example, Zhong et al. demonstrated that TREM2 binding of complement protein C1q protects against aberrant complement-mediated synaptic loss in neurodegeneration^88^. Indeed recent literature examining cases of TREM2 overexpression have found reduced inflammation and rescued cognition in AD mouse models^59,89,90^. While increasing TREM2 expression may be neuroprotective, the role of sTREM2 remains unclear. Several studies have shown affinity of sTREM2 for amyloid beta plaques in individuals with AD as well as amelioration of tau phosphorylation and associated cognitive deficits^91,92^. Findings from Moutinho et al. show that sTREM2 can modulate synaptic transmission and disrupt long-term potentiation in mouse brain suggests potentially unknown roles for sTREM2 as a neuropeptide modulator of synaptic transmission^44^. However, the induction of TREM2 and potentially sTREM2 following ethanol exposure appears to require interaction between multiple cell types and, likely, use of a human-based model system. The finding that ethanol regulates this key pathway opens a new understanding of the complex effects of alcohol ingestion on brain function.

## Methods Materials

Materials, sources, and PCR primer sequences can be found in the Supplemental Methods.

### iPSC Cell Lines

The 03SF iPSC line^93^ was used throughout with individual differentiation batches serving as replicates.

### Neural progenitor cell generation

Neural progenitor cells (NPCs) were generated as previously described^94^. Briefly, human iPSCs underwent embryoid body (EB) formation and subsequent neural induction by dual-SMAD inhibition with growth factor NOGGIN and small molecule SB431542 for 6 days. EBs were then plated on a Matrigel (Corning) coated dish in neural medium consisting of DMEF12, N2, and 1 ug/ml Laminin (Corning). Neural rosettes were manually picked and expanded in medium consisting of 1:1 DMEF12:Neurobasal, N2, B27-RA, basic fibroblast growth factor (bFGF, 20 ng/ml), SB431542 (10 µM), human leukemia inhibitory factor (hLIF, 10 ng/ml Peprotech), and CHIR99021 (3 µM, Stemcell Technologies). Expanded stocks were frozen at P3-5 for future experiments.

### Neural differentiation

To differentiate NPCs into neurons and astrocytes, NPCs were thawed and plated onto growth-factor reduced Matrigel (Corning) coated plates in neuronal differentiation medium consisting of Neurobasal (Gibco) with GlutaMAX-I supplement (Gibco), B27 supplement (1X, Gibco), Y27632 (10 µM, Reprocell), penicillin/streptomycin (P/S), Brain derived neurotrophic factor (BDNF, 10 ng/ml, PeproTech), Neurotrophin-3 (NT-3, 10 ng/mL, PeproTech), glial derived neurotrophic factor (GDNF, 10 ng/mL, PeproTech), L-Ascorbic Acid (L-AA, 200 µM, Sigma), ß-Mercaptoethanol (55 µM, Gibco). NPCs were maintained for about one week, passaged with Accutase (StemCell Technologies), and plated at 75,000 cells/cm^2^. NPC-derived cells were then maintained in neuronal maturation conditions, neuronal differentiation medium (as described above) with cAMP (1µM, Stem Cell Technologies). Half medium changes were performed every three to four days.

### Microglia progenitor cell differentiation

Microglial progenitor cells (MPCs) were generated from iPSC as previously described^17,28,29^. Briefly, human iPSCs underwent embryoid body (EB) formation and mesodermal induction by plating 20,000 cells/well in a 96-well low adherence plate (Corning) with stem cell medium (StemMACS iPSC-Brew XF; Miltenyi Biotec) supplemented with Bone Morphogenic Protein 4 (BMP4, 50 ng/ml, Peprotech), Stem Cell Factor (SCF, 20 ng/ml, Peprotech), Vascular Endothelial Growth Factor (VEGF, 50 ng/ml, Peprotech) and ß-Mercaptoethanol (55 µM; Gibco). In addition, CEPT was administered on day 0. Half medium changes were performed daily. On day 5, EBs were plated on 100 mm dishes and cultured in factory medium composed of XVIVO15 (Lonza), GlutaMAX (1X, Gibco), ß-Mercaptoethanol (55 µM, Gibco) supplemented with Macrophage Colony Stimulating factor (M-CSF, 100 ng/ml, Peprotech) and Interleukin-3 (IL-3, 25 ng/ml, Peprotech) to promote myeloid differentiation and Yolk-Sac identity of the EB. Half medium changes were performed every five days. After two to three weeks, MPCs egress into the surrounding medium and are continually produced for up to six months.

### Neuron, astrocyte, microglia triculture preparation

After seven weeks of NPC neural differentiation, microglia were added to the neuron-astrocyte culture at a density of 50,000 cells/cm^2^ and maintained for 1 week in triculture conditions (adapted from a previously published protocol^28^), with medium consisting of Neurobasal (Gibco), GlutaMAX-I (Gibco), B27 (Gibco), P/S, supplemented with BDNF (10 ng/ml, PeproTech), NT-3 (10 ng/mL, PeproTech), GDNF (10 ng/mL, PeproTech), L-Ascorbic Acid (L-AA, 200 µM, Sigma), ß-Mercaptoethanol (55 µM, Gibco), M-CSF (25 ng/ml, Peprotech), Interleukin-34 (IL-34, 100 ng/ml, Peprotech), and granulocyte-macrophage colony-stimulating factor (GM-CSF, 10 ng/ml, Peprotech). After one week, M-CSF was removed from the culture medium.

### Intermittent Ethanol Exposure (IEE)

After eight weeks of culture, a 7-day intermittent ethanol exposure was performed as previously described^30^. Briefly, fresh medium consisting of Neurobasal (Gibco), GlutaMAX-I (Gibco), B27 (Gibco), P/S, supplemented with BDNF (10 ng/ml, PeproTech), NT-3 (10 ng/mL, PeproTech), GDNF (10 ng/mL, PeproTech), L-Ascorbic Acid (L-AA, 200 µM, Sigma), ß-Mercaptoethanol (55 µM, Gibco), Interleukin-34 (IL-34, 100 ng/ml, Peprotech), and granulocyte-macrophage colony-stimulating factor (GM-CSF, 10 ng/ml, Peprotech) was mixed with conditioned medium at a 3:1 (v/v) ratio to provide a 75% medium change. On the first day of IEE treatment, all medium was removed from each well and replaced with the conditioned medium mix supplemented with ethanol (200 Proof, Decon Laboratories) to give final concentrations of 0 mM, 20 mM, 40 mM, or 75 mM. On days two and three, half the medium was removed from each well, ethanol was added at twice the required concentrations, and the mix was added back to the cultures. On day four, a half medium change was performed with fresh medium also supplemented with twice the required concentrations of ethanol for each condition. Days five and six were the same as days two and three. On day seven, cultures were fixed for immunocytochemistry and RNA isolation.

### Immunocytochemistry

Immunocytochemistry was performed as previously described ^17,30,95^. Cells were fixed in ice-cold 4% PFA, PBS for 15 min at room temperature. Following fixation, samples were permeabilized with 0.2% Triton X-100, PBS for 7 minutes, and incubated with blocking buffer (5% natural goat serum, 4% bovine serum albumin, and 0.05% triton X-100, PBS) for 45 minutes. Primary antibodies were diluted in blocking buffer and incubated either at room temperature for 1 hour or overnight at 4°C. Following primary incubation, five washes were performed with PBS, 0.05% Triton (PBST). Secondary antibodies diluted in blocking buffer (1:500 dilution) were then incubated at room temperature for 1 hour. Following secondary incubation, five washes were performed with PBST. Finally, coverslips were mounted onto glass slides with Fluoroshield with DAPI (Sigma). The coverslip was then sealed to the slide with clear nail polish.

### Image Acquisition and Processing

15 images were taken per condition using a Zeiss LSM700 Confocal Microscope at 20X magnification. To enable blinding analysis, the ImageJ plugin Filerandomizer was used to assign arbitrary file names to each file.

#### Region of interest generation

For each image, regions of interest (ROIs) were generated around individual microglia using ImageJ. To do so, the IBA1 channel was isolated, auto adjusted for brightness/contrast, auto-thresholded (triangle setting), and analyzed with the Analyze Particle plugin. When needed, ROIs were manually corrected after auto-generation using the Selection Brush tool on ImageJ. Careful comparison with the original image was maintained throughout manual adjustments. Incomplete cells at the edge of the image and significantly overlapping/ambiguous cells were excluded from analysis.

#### Fluorescence Quantification

To quantify fluorescence, ROIs from the IBA1 masks were overlayed on the CD68 channel. The integrated intensity, area, and perimeter were then measured for each ROI using ImageJ. To properly quantify CD68 expression, the corrected total cell fluorescence was calculated according to the formula: CTCF = Integrated Density – (Area of selected cell X Mean fluorescence of background readings).

#### FracLac for ImageJ

For detailed multiparameter analysis of morphology (e.g. fractal dimension, lacunarity, convex hull) the free online software *FracLac for ImageJ* was used^37^. Batch analysis was performed on binary image outlines generated from the IBA1 masks for streamlined processing.

### RNA Purification

To isolate RNA, a Qiagen RNeasy Plus Mini Kit (cat. #74136) was used. Approximately1.5 – 2 million cells were lysed in RLT Plus lysis buffer. The Lysate was then homogenized with a QIAshredder and centrifuged for 2 minutes at 15,000 RPM. The homogenized lysate was transferred to a gDNA eliminator spin column placed in a 2 mL collection tube and centrifuged for 30 seconds at 15,000 RPM to eliminate genomic DNA. One volume of 70% ethanol was added to the flow-through and mixed. The sample was then transferred to a RNeasy Mini spin column in a 2mL collection tube and centrifuged for 30 seconds at 15,000 RPM. Buffer RW1 was added to the RNeasy Mini spin column and centrifuged followed by Buffer RPE. Total RNA was then eluted using RNase-free water and quantified with a NanoDrop Spectrophotometer (Thermo Scientific).

### RNA Sequencing

RNA samples were submitted to Novogene for library preparation and sequencing. Data were processed through a standard workflow at Novogene for genome alignment, gene mapping, and quantification. Gene-level analysis was performed starting with a raw counts matrix using edgeR^96^ and a generalized linear model in R/Bioconductor. For isoform analysis, fastq files were mapped to a reference library of Ensembl cDNA transcripts with Kallisto^97^, then scaled and normalized in DESeq2. Results are available from the NIH Gene Expression Omnibus archive (Accession GSE303551).

### RT-qPCR

cDNA was synthesized using approximately 75-100 ng of RNA using the SuperScript VILO cDNA synthesis Kit (Invitrogen, cat. #11755050). RT-qPCR using PowerUp SYBR Green Master Mix (ThermoFisher; cat. #A25743) was then performed. RT-qPCR reactions were performed in QuantStudio 3 system (ThermoFisher) using the following parameters: 50°C for 2 minutes and 95°C for 2 minutes, then 95°C for 15 seconds and 60°C for 1 minute for 40 cycles. Primers were designed using NIH Primer-BLAST. Duplicate reactions were performed in five biological replicates.

### Statistical Analysis

All data are presented as the mean ± standard error of the mean (SEM), unless otherwise indicated. All experiments were repeated at least three times unless otherwise noted. The effects of ethanol on treated groups versus untreated controls, as well as between-condition comparisons, were compared using one-way ANOVA followed by post-hoc significance tests. To indicate statistical significance, the following symbols were used *p < 0.05, **p < 0.01, ***p < 0.001, ****p < 0.001.

## Author Contributions

AJB, ZPP and RPH conceived the study. AJB and YA designed experiments and interpreted data. AJB and YA performed most of the experiments with technical assistance from XL and ACS. AJB, ACS, and YA performed RT-qPCR. AJB and YA performed immunocytochemistry, immunohistochemistry, and confocal imaging. AJB, RPH, SZ, XL, and JD performed computational analysis of RNAseq data. AJB prepared the figures and wrote the manuscript with input from all co-authors.

## Acknowledgements

This study was supported by grants NIAAA R01AA023797 and U10AA008401. The Child Health Institute of New Jersey is supported in part by the Robert Wood Johnson Foundation (RWJF grant #74260). JD was supported by R01MH106575, R01AG063175 and R01AG081374. AJB was supported by NIGMS NIH T32GM008339 and by NCATS NIH TL1TR003019.

## MAIN FIGURE TITLES AND LEGENDS

**Supplemental Figure 1:**
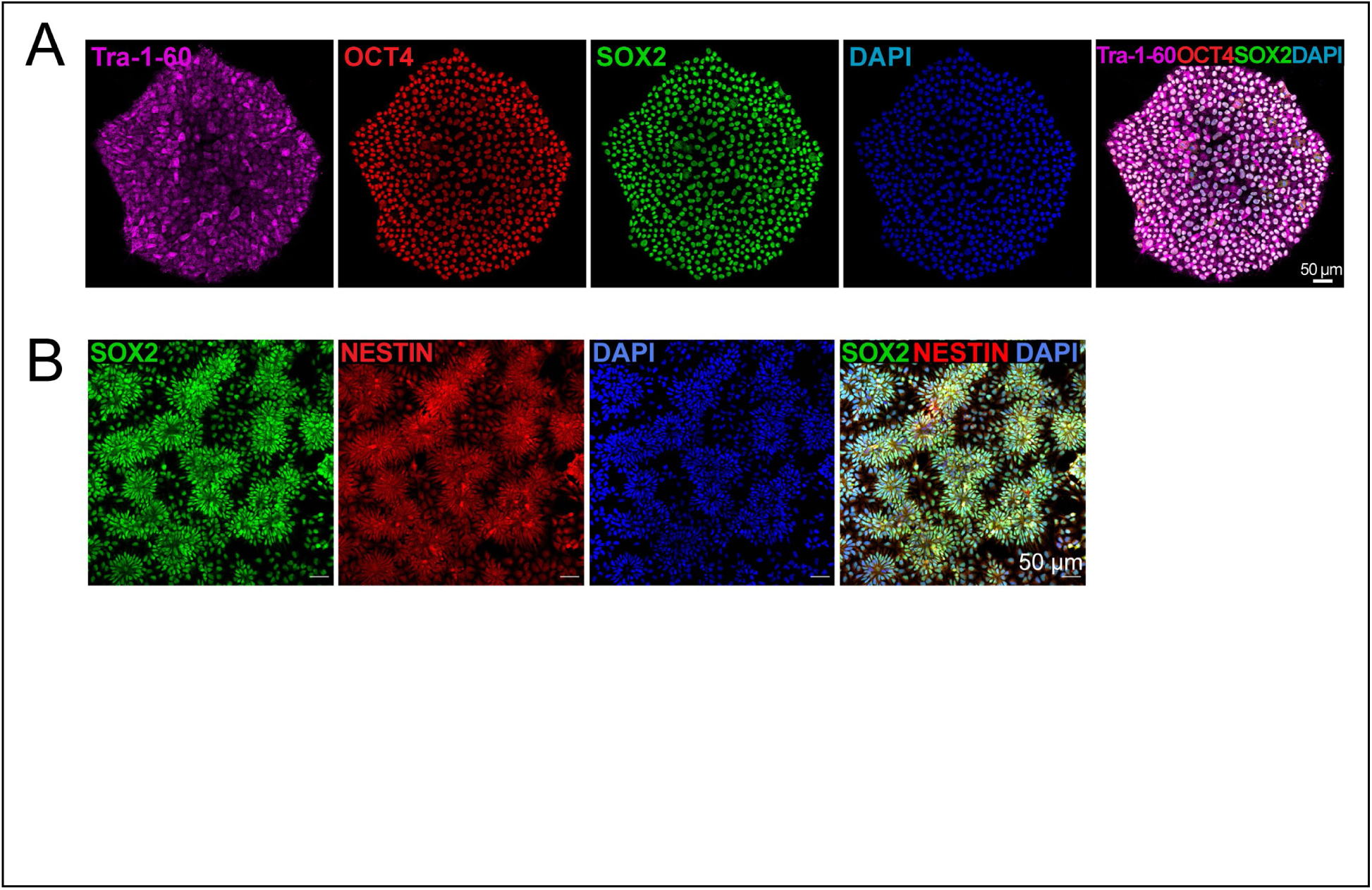
Characterization of 03SF cell line pluripotency and differentiation. (A) Representative confocal images of immunocytochemical staining against pluripotency markers TRA-1-60, OCT-4, SOX2, and DAPI nuclei. (B) Representative confocal images of staining against neural progenitor cell markers SOX2, NESTIN, and DAPI nuclei.

**Supplemental Figure 2:**
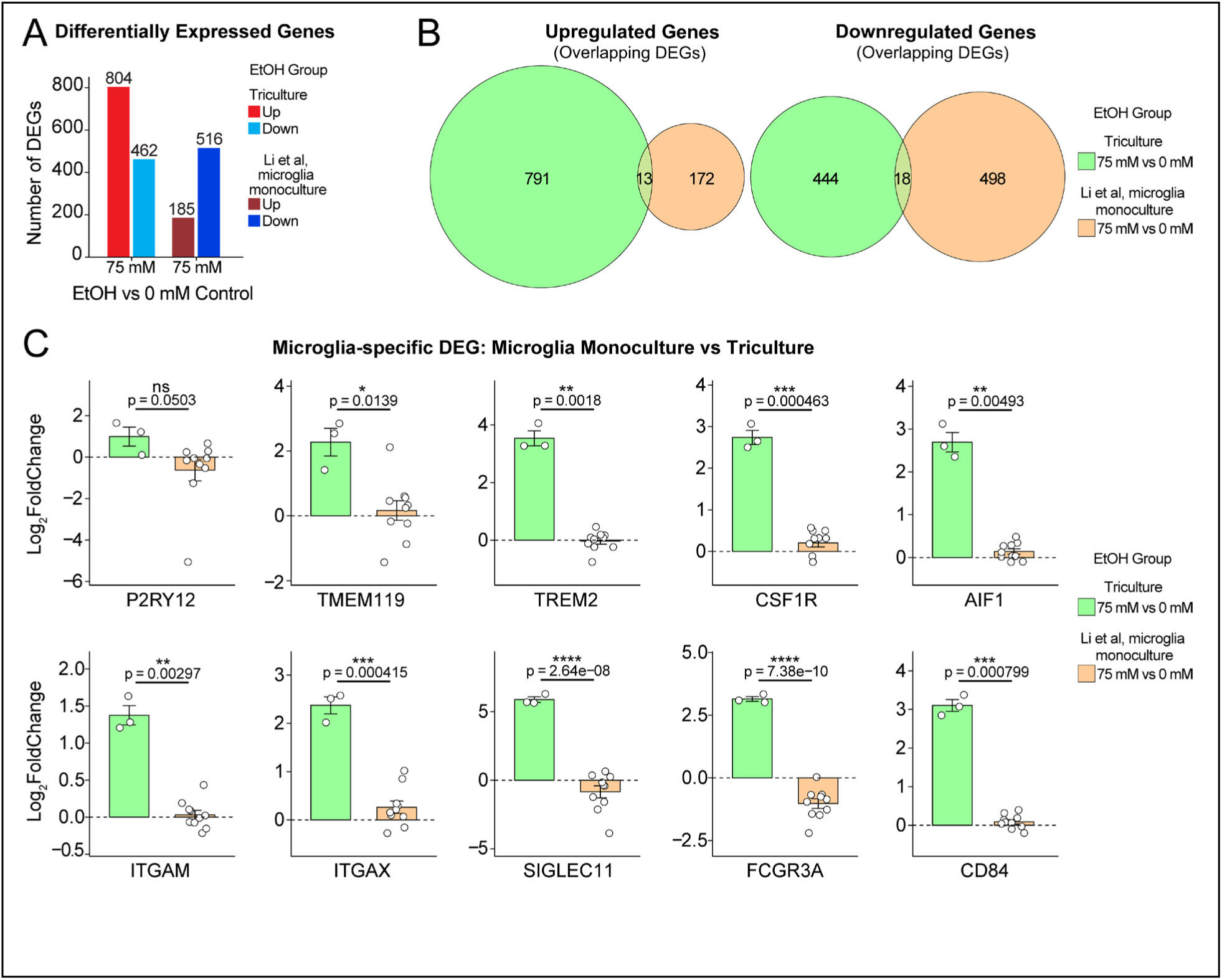
DEG comparison of triculture vs microglia monoculture. (A) A barplot showing the total number of upregulated and downregulated differentially expressed genes (DEG) for all three triculture conditions and microglia monoculture 75 mM vs 0mM ethanol from Li et al, 2025^18^. (B) Venn diagrams showing numbers of overlapping and unique DEG between the various conditions from Panel A. (C) A barplot of the Log_2_FoldChange of microglia-specific genes, comparing 75 mM vs 0 mM intermittent ethanol exposure for both the triculture 75 mM condition and the microglia monoculture 75 mM condition. *p < 0.05, **p < 0.01, ***p <0.001, ****p < 0.0001. Data are presented as means ± SEM.

## Supplemental Methods

### Source of reagents

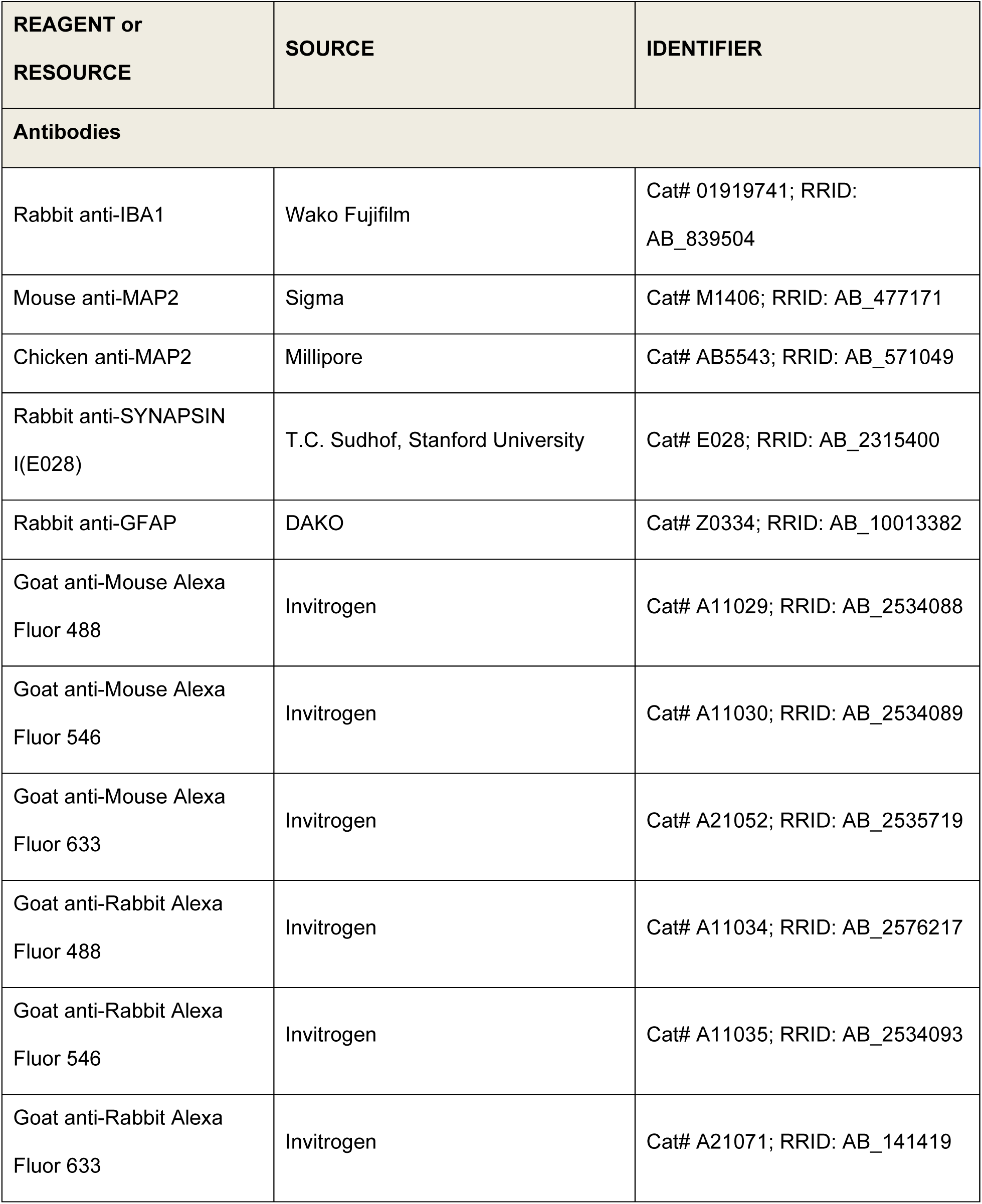
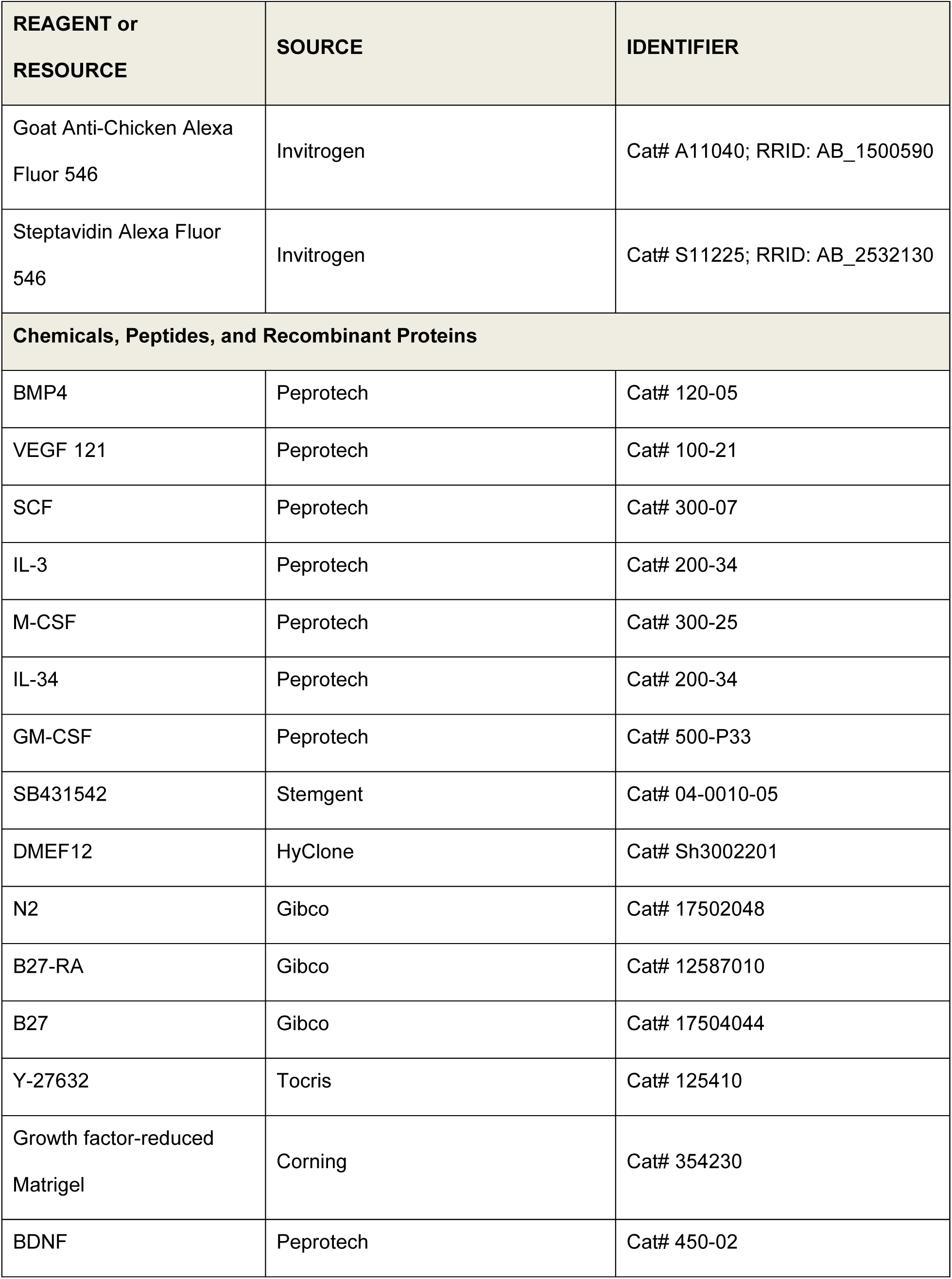
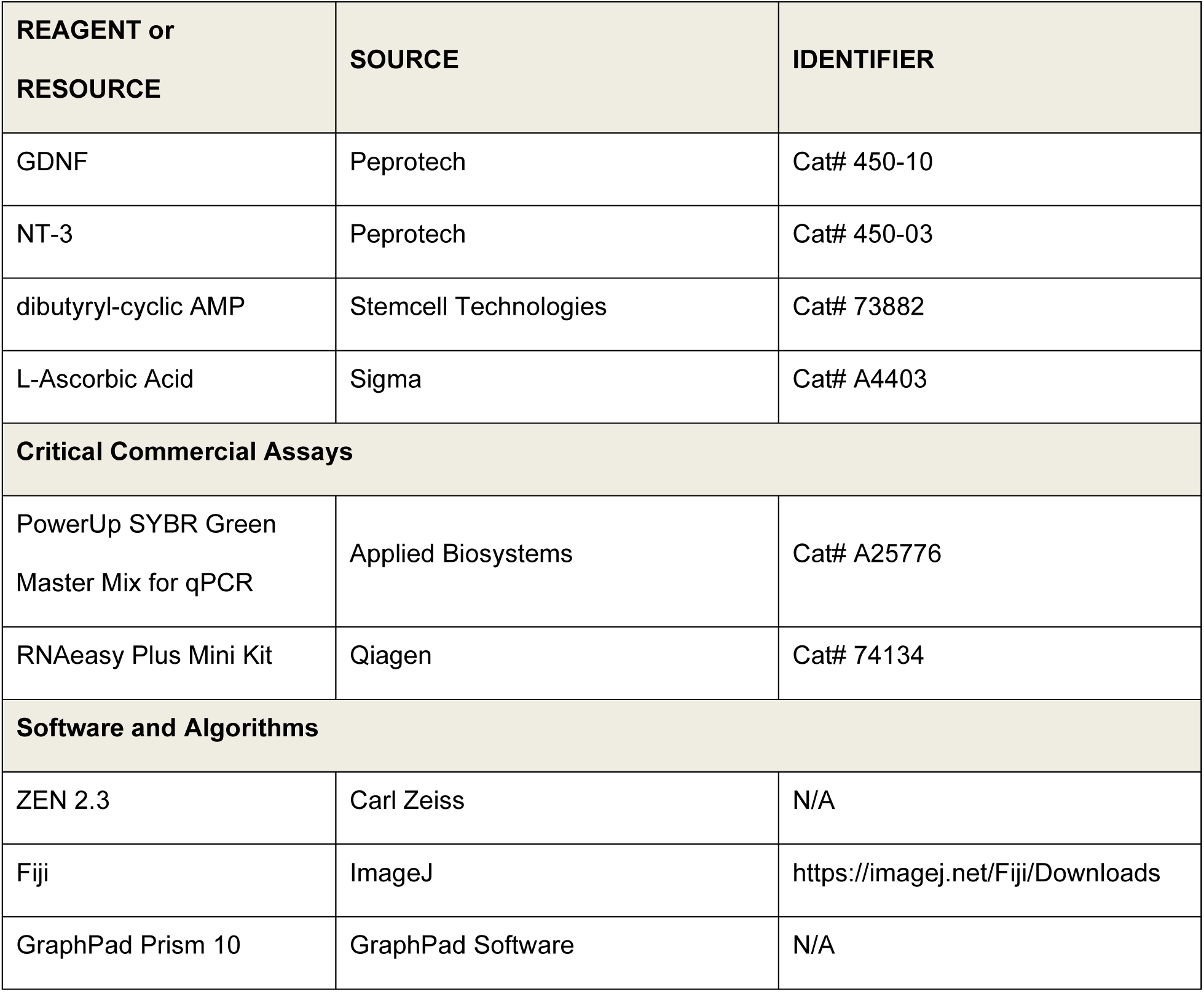

| REAGENT or RESOURCE | SOURCE | IDENTIFIER |
| --- | --- | --- |
| **Antibodies** |  |  |
| Rabbit anti-IBA1 | Wako Fujifilm | Cat# 01919741; RRID: AB_839504 |
| Mouse anti-MAP2 | Sigma | Cat# M1406; RRID: AB_477171 |
| Chicken anti-MAP2 | Millipore | Cat# AB5543; RRID: AB_571049 |
| Rabbit anti-SYNAPSIN I(E028) | T.C. Sudhof, Stanford University | Cat# E028; RRID: AB_2315400 |
| Rabbit anti-GFAP | DAKO | Cat# Z0334; RRID: AB_10013382 |
| Goat anti-Mouse Alexa Fluor 488 | Invitrogen | Cat# A11029; RRID: AB_2534088 |
| Goat anti-Mouse Alexa Fluor 546 | Invitrogen | Cat# A11030; RRID: AB_2534089 |
| Goat anti-Mouse Alexa Fluor 633 | Invitrogen | Cat# A21052; RRID: AB_2535719 |
| Goat anti-Rabbit Alexa Fluor 488 | Invitrogen | Cat# A11034; RRID: AB_2576217 |
| Goat anti-Rabbit Alexa Fluor 546 | Invitrogen | Cat# A11035; RRID: AB_2534093 |
| Goat anti-Rabbit Alexa Fluor 633 | Invitrogen | Cat# A21071; RRID: AB_141419 |
| Goat Anti-Chicken Alexa Fluor 546 | Invitrogen | Cat# A11040; RRID: AB_1500590 |
| Streptavidin Alexa Fluor 546 | Invitrogen | Cat# S11225; RRID: AB_2532130 |
| **Chemicals, Peptides, and Recombinant Proteins** |  |  |
| BMP4 | Peprotech | Cat# 120-05 |
| VEGF 121 | Peprotech | Cat# 100-21 |
| SCF | Peprotech | Cat# 300-07 |
| IL-3 | Peprotech | Cat# 200-34 |
| M-CSF | Peprotech | Cat# 300-25 |
| IL-34 | Peprotech | Cat# 200-34 |
| GM-CSF | Peprotech | Cat# 500-P33 |
| SB431542 | Stemgent | Cat# 04-0010-05 |
| DMEF12 | HyClone | Cat# Sh3002201 |
| N2 | Gibco | Cat# 17502048 |
| B27-RA | Gibco | Cat# 12587010 |
| B27 | Gibco | Cat# 17504044 |
| Y-27632 | Tocris | Cat# 125410 |
| Growth factor-reduced Matrigel | Corning | Cat# 354230 |
| BDNF | Peprotech | Cat# 450-02 |

| REAGENT or<br>RESOURCE | SOURCE | IDENTIFIER |
| --- | --- | --- |
| GDNF | Peprtech | Cat# 450-10 |
| NT-3 | Peprtech | Cat# 450-03 |
| dibutyryl-cyclic AMP | Stemcell Technologies | Cat# 73882 |
| L-Ascorbic Acid | Sigma | Cat# A4403 |
| **Critical Commercial Assays** |  |  |
| PowerUp SYBR Green<br>Master Mix for qPCR | Applied Biosystems | Cat# A25776 |
| RNAeasy Plus Mini Kit | Qiagen | Cat# 74134 |
| **Software and Algorithms** |  |  |
| ZEN 2.3 | Carl Zeiss | N/A |
| Fiji | ImageJ | https://imagej.net/Fiji/Downloads |
| GraphPad Prism 10 | GraphPad Software | N/A |

### Oligonucleotide primer sequences

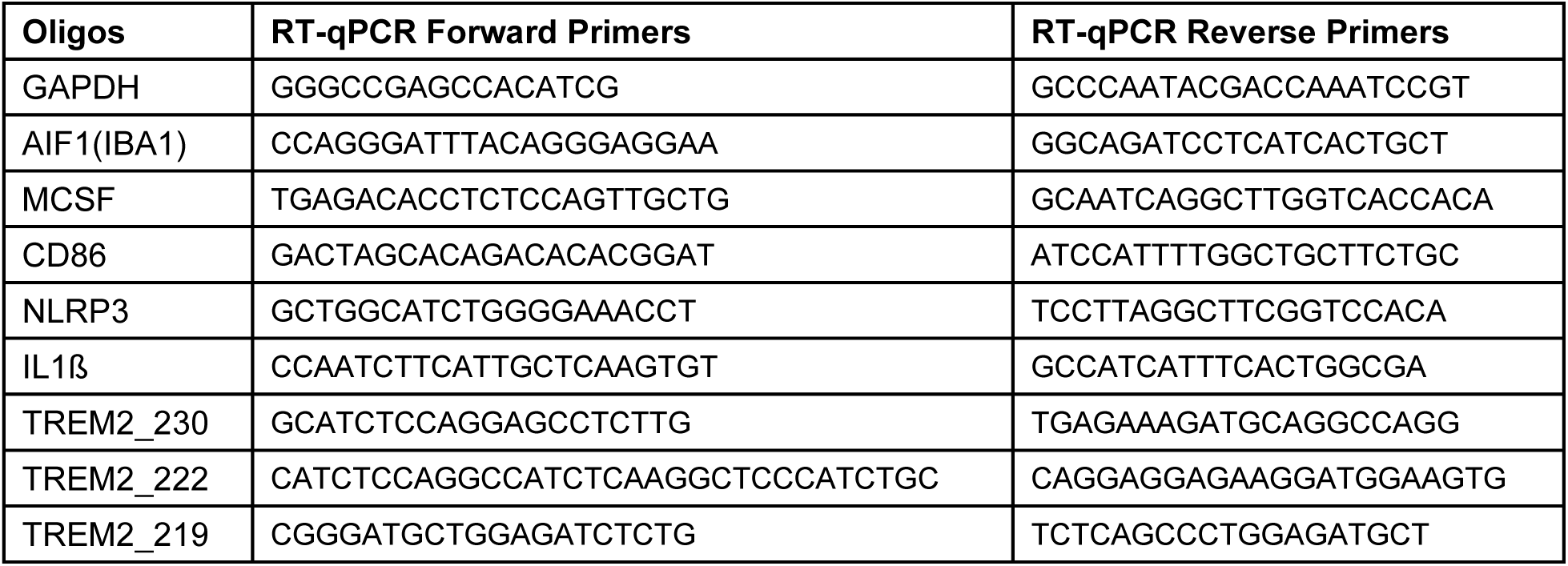

